# *Microcystis*-triggered shifts in the symbiotic microbiome of *Myriophyllum spicatum* rapidly suppress *Microcystis aeruginosa*

**DOI:** 10.64898/2026.08.27.747664

**Authors:** Seonah Jeong, Hayoung Lee, So-Ra Ko, Dong-Yun Choi, Won-Suk Choi, Yuna Shin, Kyunghyun Kim, Hee-Sik Kim, Chi-Yong Ahn

**Affiliations:** Cell Factory Research Center, Korea Research Institute of Bioscience & Biotechnology (KRIBB), 125 Gwahak-ro, Yuseong-gu, Daejeon, 34141, Republic of Korea; Department of Environmental Biotechnology, KRIBB School of Biotechnology, University of Science and Technology, Daejeon, 34113, Republic of Korea; Han River Environmental Research Center, Water Environmental Research Department, National Institute of Environmental Research, Incheon 22689, Republic of Korea

**Keywords:** *Microcystis aeruginosa*, Cyanobacterial harmful algal blooms, Bloom control, Microbiome, Submerged macrophyte, *Myriophyllum spicatum*, Meta-transcriptomics

## Abstract

While the suppression of toxic cyanobacteria by aquatic plants has long been recognized, few studies have clearly differentiated between the allelopathic effects of the plant itself and the inhibitory influence of its associated microbiome. This study aimed to clarify the primary inhibitory agent by pre-culturing *Myriophyllum spicatum* (Eurasian watermilfoil) under three conditions: (1) BG11 medium, (2) live *Microcystis aeruginosa* KW culture, and (3) a *Microcystis*-symbiotic microbiome (excluding *Microcystis* cells). After a 7-day pre-culture, *Myriophyllum* shoots were transferred to fresh *Microcystis* culture. The *Myriophyllum* pre-cultured in *Microcystis* culture exhibited rapid inhibition against *Microcystis* (84% within day 1), whereas the *Myriophyllum* pre-cultured in BG11 medium showed delayed responses (89% by day 7). In contrast, inhibition remained below 50% in the *Myriophyllum* pre-cultured with the *Microcystis*-symbiotic microbiome. Notably, plant-derived soluble compounds exhibited weak inhibitory effects, whereas the microbiome showed stronger inhibitory activity, indicating that the plant-associated microbiome plays a more dominant role than the plant itself. Exposure to *Microcystis* triggered significant shifts in plant-symbiotic microbial community composition, leading to rapid enhancement of inhibitory activity in the *Myriophyllum* microbiome. Microbial community analysis identified 28 bacterial taxa closely associated with the inhibitory response, including strains involved in organic matter degradation, adhesion, biofilm formation, and predatory behavior. Meta-transcriptomic analysis further confirmed increased expression of genes related to bacterial adhesion, biofilm formation, and carbohydrate metabolism following *Microcystis* exposure, highlighting functional adaptations linked to cyanobacterial suppression. These findings underline the role of microbiome-mediated cyanobactericidal mechanisms, providing new insights into a nature-based solution for mitigating *Microcystis*-dominated harmful algal blooms.

## 1. Introduction

The increasing occurrence of cyanobacterial harmful algal blooms (cyanoHABs) has become a pressing global environmental concern, closely linked to the multifaceted impacts of climate change. Over the past few decades, the frequency, intensity, and geographic expansion of cyanoHABs have risen significantly, posing serious risks to aquatic ecosystems and public health (Chorus and Welker, 2021). These blooms, primarily driven by eutrophication in freshwater systems, lead to water quality deterioration, ecological imbalances, and the production of toxins that endanger both biodiversity and human health. *Microcystis aeruginosa*, the most famous bloom-forming cyanobacterium, has been the predominant causative agent in freshwater environments such as lakes, ponds, and rivers since the 1980s (Feng et al., 2024; Huisman et al., 2018; Paerl and Barnard, 2020). Toxic *Microcystis* cells often form aggregates with other freshwater bacteria, facilitating metabolite exchange and cooperative defense mechanisms (Hoke et al., 2021; Xie et al., 2016).

Various mitigation strategies have been developed to address cyanoHABs, including nutrient management, physical and chemical treatments, and biological control (Kibuye et al., 2021a; Kibuye et al., 2021b). Nutrient management focuses on reducing nitrogen and phosphorus inputs from agricultural and urban sources through advanced land-use practices and wastewater treatment technologies (Igwaran et al., 2024). Despite progress in these methods, significant challenges remain, such as high costs, secondary pollution, and the limited scalability of current strategies (Kibuye et al., 2021a; Kibuye et al., 2021b; Meyer et al., 2017). Furthermore, these approaches often fail to consider the complex interactions among aquatic organisms, and the broader ecosystem. Understanding and leveraging these interactions could facilitate the development of more effective and sustainable solutions for cyanoHAB management, particularly in the face of ongoing climate change and increasing anthropogenic pressures.

Biological control methods, including the use of bacteria and aquatic organisms with antagonistic effects on cyanobacteria, have attracted significant research interest for their potential in sustainable cyanoHAB management (Coyne et al., 2022; Kang et al., 2024). Among these strategies, aquatic plants with water purification capabilities have shown considerable promise. Through nutrient uptake and allelopathic interactions, these plants can inhibit cyanobacterial growth while minimizing ecological disturbances (Nezbrytska et al., 2022; Wang and Liu, 2023). Recent studies highlight the critical role of symbiotic microorganisms associated with aquatic plants in environmentally friendly water quality management (Chen et al., 2023; Imai et al., 2021; Jiang et al., 2019; Miyashita, 2019). For instance, *Pseudomonas* sp. Go58 isolated from aquatic plant *Trapa jeholensis*, has been found to secrete chemical compounds that disrupt cyanobacterial cell integrity (Chen et al., 2023). Symbiotic microorganisms contribute to ecological balance by influencing nutrient cycling, synthesizing bioactive compounds, and mitigating cyanoHABs (Park et al., 2024; Cai et al., 2024).

Among submerged macrophytes, *Myriophyllum spicatum* (Eurasian watermilfoil) is particularly notable for its ability to control cyanoHABs. This plant efficiently absorbs nitrogen and phosphorus, reducing nutrient availability and thereby limiting cyanobacterial proliferation (Bao et al., 2020). Additionally, *Myriophyllum* releases allelochemicals that inhibit cyanobacterial growth (Jeong et al., 2021; Leu et al., 2002; Mohamed et al., 2017). These mechanisms underscore the ecological importance of *Myriophyllum* in freshwater ecosystems and highlight its potential for nature-based water management practices. Despite extensive research on macrophyte-cyanobacteria interactions, the role of symbiotic microbiomes in cyanobacterial suppression remains largely unexplored.

This study examined how the *Myriophyllum spicatum*-associated microbiome contributes to *Microcystis aeruginosa* suppression, distinguishing microbial influence from plant allelopathic effects. Additionally, we analyzed microbial community shifts and functional adaptations involved in this process. By exploring the interactions between the *Myriophyllum*-symbiotic microbiome and *Microcystis*, this research provides novel insights into the microbial mechanisms driving cyanobacterial suppression and underscores the potential of submerged aquatic macrophytes as a nature-based solution (NbS) for managing *Microcystis* blooms in freshwater ecosystems.

## 2. Materials and Methods

### 2.1. Pre-culture and co-culture experimental design

*Myriophyllum spicatum* (Ms) plants used in this study were collected from the Yudeung Stream (36° 17’ 11.4” N, 127° 22’ 39.2” E) in Daejeon, South Korea. The collected plants were rinsed three times with on-site water to remove surface debris.

For pre-culture phase, 5-cm shoot segments were cultured for 7 days in 125 mL under three different conditions: (1) Group Pre-BG11, Ms pre-cultured in BG11 medium; (2) Group Pre-Ma, Ms pre-cultured in *Microcystis aeruginosa* KW (Ma) culture (8.7 × 10^5^ cells/mL); and (3) Group Pre-Sym, Ms pre-cultured in the *Microcystis*-symbiotic microbiome solution (Ma excluded by filtering through a 0.8-μm membrane filter, BG11 diluted to 10%). During the pre-culture, active chlorophyll concentrations were measured on days 0, 2, 4, 6, and 7 using a PHYTO-PAM phytoplankton analyzer (Walz, Effeltrich, Germany).

*Microcystis aeruginosa* KW was isolated from the Wangsong Reservoir in South Korea (Srivastava et al., 2016 and 2019). Ma culture was grown in sterile BG-11 medium at 25 °C under constant light (80 μmol photons/m^2^/s). Ma was grown predominantly as single cells in flask culture condition.

For co-culture experiments, the Ms shoots from each pre-culture group were transferred into 125 ml of fresh Ma culture (8.7 × 10^5^ cells/mL), after 7 days of pre-culture under the respective condition (Figs. 1 and S1). Three biological replicates were initially prepared; however, one replicate was lost during the experiment. Throughout the co-culture of Ms and Ma, active chlorophyll concentrations of the treatment and control groups were measured on days 0, 1, 3, 5, 7, and 9.

**Figure 1.**
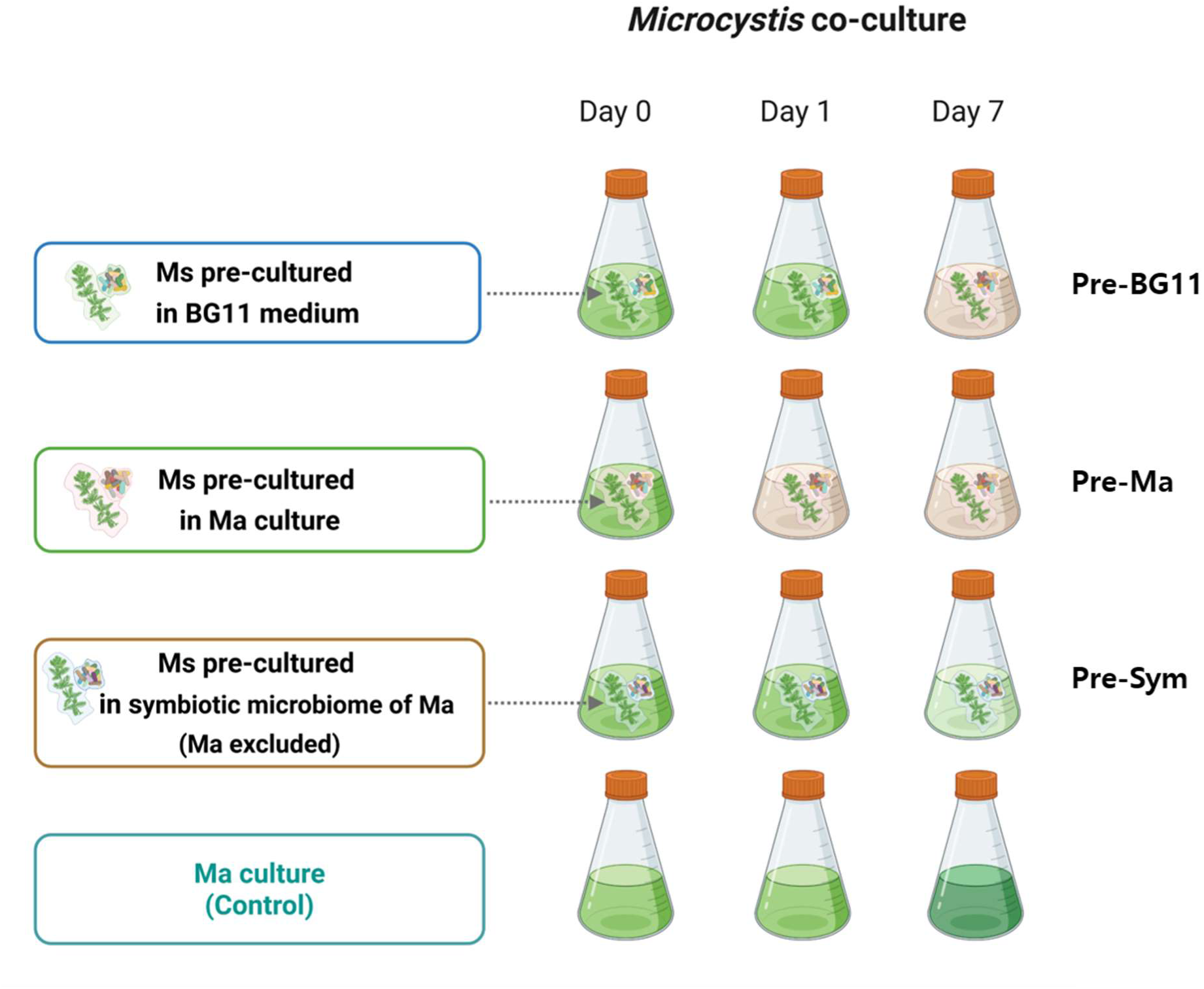
Experimental design for investigating *Microcystis*-triggered inhibitory responses of *Myriophyllum spicatum* (Ms) against *Microcystis aeruginosa* (Ma). The pre-culture phase consisted of three distinct conditions. After 7 days of pre-culture, co-culture experiments were conducted by transferring Ms shoots from each condition into fresh Ma culture.

### 2.2. Calculation of growth inhibition

Growth inhibition (*I*) of *Microcystis* was calculated using the following equations (Jeong et al., 2021):

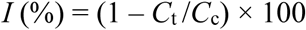

where *C*_t_ and *C*_c_ values are active chlorophyll concentrations of the treatment and control, respectively.

### 2.3. Comparison of inhibitory effects of plant-derived chemicals and plant-associated microbiome

Ms plants cultivated in a pond at KRIBB (36° 22’ 36.6” N, 127° 21’ 30.5” E) were used in this experiment. A 10-cm shoot segment was excised and gently rinsed with pond water to remove surface debris. The shoot was then briefly rinsed with distilled water to remove any remaining debris. The plant segment was then trimmed to a 5-cm shoot (0.45 g wet weight) and co-cultured with Ma (250 ml, 8.5 × 10^5^ cells/mL) for 12 days, until active chlorophyll levels of the cyanobacteria decreased significantly.

Once the growth of Ma was inhibited, the culture solution was divided into three experimental groups (Fig. 2A), which were designed to distinguish the relative contributions of dissolved plant-derived compounds and the plant-associated microbiome to the inhibition of Ma. Group F was used to evaluate the effect of dissolved compounds after removal of microbial cells. Group S represented the whole co-culture-derived fraction containing both dissolved compounds and the plant-associated microbiome. Group M was used to assess whether the microbiome detached from the plant surface could inhibit Ma growth. This design allowed us to compare chemical-only, both chemical-/microbiome-containing, and microbiome-only treatments under the same Ma inoculation conditions.

**Figure 2.**
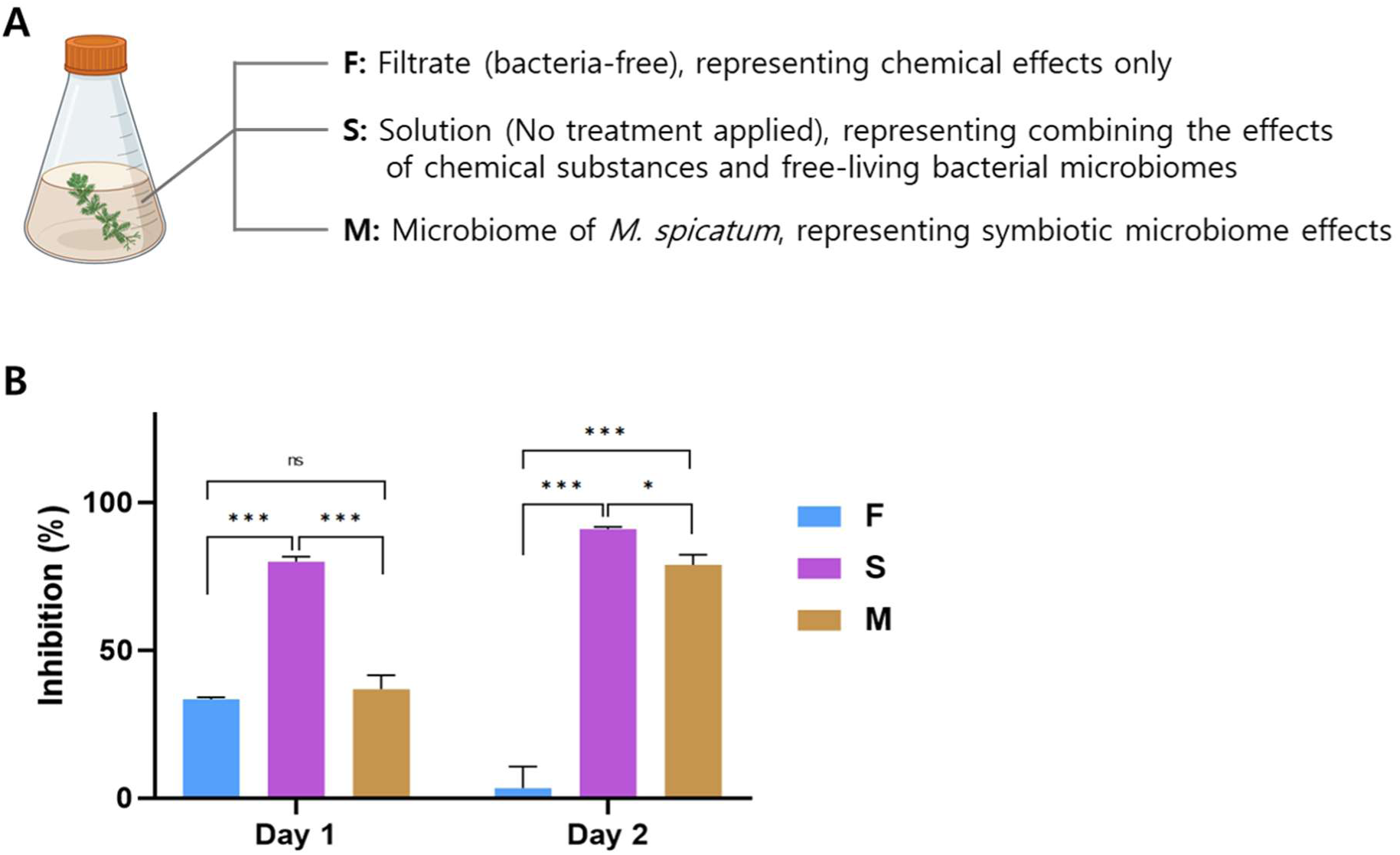
**A**, schematic representation of the three treatment conditions (F, S, and M) derived from the co-culture of Ma and Ms. **B**, inhibition of Ma by three treatment groups based on active chlorophyll (mean ± SD). Significant differences were observed among groups on both experimental days using one-way ANOVA (day 1, *p* = 1.97 × 10^-6^; day 2, *p* = 8.69 × 10^-7^). Pairwise comparisons were conducted using Tukey’s HSD post-hoc test; *, *p* < 0.05; **, *p* < 0.01; ***, *p* < 0.001; ns, not significant.

Each experimental group was prepared as follow: (1) Group F, the filtrate obtained by passing the culture using a 0.22-µm membrane filter (Advantec MFS, Inc., Dublin, CA, USA), thereby removing the plant-associated microbiome while retaining plant-derived chemicals; (2) Group S, the unfiltered culture containing both plant-associated microbiome and plant-derived chemicals; and (3) Group M, the microbiome detached from the plant shoot and resuspended in an equal volume (250 ml) of fresh BG11 medium, thereby excluding plant-derived chemicals. Each experimental group was inoculated with Ma at 10% (v/v), resulting in a final concentration of 8.5 × 10^5^ cells/mL in a total volume of 30 mL. The experiment was performed in biological triplicate for 2 days. Active-chlorophyll was measured daily, and Ma cell counting was conducted on day 2.

### 2.4. DNA extraction, PCR amplification, and amplicon sequencing

For microbial community analysis, 10 mL of each sample was filtered on day 1, 3 and 7 using a 0.22-μm mixed cellulose membrane filter (Advantec MFS, Inc., Dublin, CA, USA). Genomic DNA was extracted from the filtered samples stored at −80°C. DNA extraction was performed using a ChargeSwitch® Forensic DNA Purification Kit (Thermo Fisher Scientific, Waltham, MA, USA). The V3–V4 region of the 16S rRNA gene was amplified using the universal bacterial primer set 341F/805R (341F: CCTACGGGNGGCWGCAG; 805R: GACTACHVGGGTATCTAATCC) (Herlemann et al., 2011). PCR amplification was carried out using Ex Taq™ Hot Start Version (Takara Bio Inc., Shiga, Japan). The resulting PCR products were purified using Agencourt AMPure XP beads (Beckman Coulter, Brea, CA, USA). DNA concentration was quantified using a Quant-iT dsDNA HS Assay Kit (Thermo Fisher Scientific), and the purified amplicons were pooled in equimolar concentrations. Paired-end sequencing was conducted on the samples using an Illumina MiSeq (Illumina, San Diego, CA, USA) by Macrogen Corporation (Seoul, South Korea), generating 2 × 300 bp reads.

### 2.5. Microbial community structure analysis

Raw sequences were processed using DADA2, following the amplicon sequence variant (ASV) standard operating procedure (Callahan et al., 2016). Low-quality and ambiguous sequences were removed, along with sequences containing more than two mismatches to the forward primer, more than one mismatch to the barcode, or lengths outside the range of 200 – 270 bp. The Silva database v138.1 was used for sequence alignment and classification (Quast et al., 2012). Non-bacterial ASVs, including those of Archaea, as well as chloroplasts and mitochondria from Eukaryota, were removed. In addition, rare ASVs (singletons, doubletons, and tripletons) were excluded from further analysis. The resulting dataset was normalized, and ASVs were ordered based on their relative abundances in descending order.

### 2.6. Identification of key bacteria related to *Microcystis* inhibition

Random Forest regression was employed to identify key bacterial taxa associated with Ma inhibition, due to its capability to handle complex, non-linear relationships in microbial community data (Wassan et al., 2019). The samples were labeled into two classes: I (inhibition over 50%) and N (inhibition below 50%). For data preprocessing, a variance filter (interquartile range 25%) and Pareto scaling were applied. Multivariate receiver operating characteristic (ROC) curve analysis was conducted to automate important feature identification and evaluate model performance. ROC curves were generated using Monte-Carlo cross validation (MCCV) with balanced sub-sampling. In each MCCV, two-thirds of the samples were used to evaluate feature importance. The top-ranked features (up to 100) were then utilized to build classification models, which were validated on the remaining one-third of the samples. This process was repeated multiple times to calculate the performance metrics and confidence interval for each model.

### 2.7. Meta-transcriptome analysis

Metatranscriptomic analysis was performed to examine the transcriptional responses of the microbial community associated with the Ms-Ma co-culture at the time when *Microcystis* growth inhibition became evident. Samples from the control and Ms-treated groups were collected on day 7 of co-culture, filtered through 0.22-μm mixed cellulose membrane filter, preserved with 1 mL of RNA*later*^TM^ (Invitrogen, Carlsbad, CA, USA). All filters were then stored in cryovials at −80°C until DNA and RNA extraction.

Total RNA was quantified using Quant-iT RiboGreen (Invitrogen, Carlsbad, USA) and integrity was assessed using the TapeStation RNA ScreenTape (Agilent Technologies, Santa Clara, USA). Libraries were prepared from 0.5 μg of total RNA using the Illumina Stranded Total RNA Library Prep with Ribo-Zero Plus Microbiome kit (Illumina, San Diego, USA). Following rRNA depletion, RNA was fragmented and reverse-transcribed into cDNA, followed by second-strand synthesis, adapter ligation, and PCR amplification. Libraries were quantified and quality-checked, then sequenced on an Illumina NovaSeqX platform (2×150 bp). The obtained sequencing data were analyzed using SAMSA2, an upgraded version of Simple Annotation of Metatranscriptomes by Sequence Analysis (SAMSA) (Westreich et al., 2018).

## 3. Results

### 3.1. Enhanced inhibition of *Microcystis* following pre-exposure

During the pre-culture phase, the active chlorophyll concentration in the “Ma + Ms” co-culture began to decline from day 4 compared to the control group (Fig. 3). By day 7, the active chlorophyll concentration in the control group reached 257 μg/L, whereas the “Ma + Ms” treatment decreased markedly to 46 μg/L. In the subsequent co-culture experiment, Group Pre-Ma demonstrated a rapid onset of inhibitory effects on *Microcystis*, achieving 84% inhibition within the first day (Figs. 1 and 4). Group Pre-BG11 exhibited a slower inhibitory response, reaching 89% inhibition by day 7. However, inhibition in the Group Pre-Sym remained below 50% throughout the experimental period.

**Figure 3.**
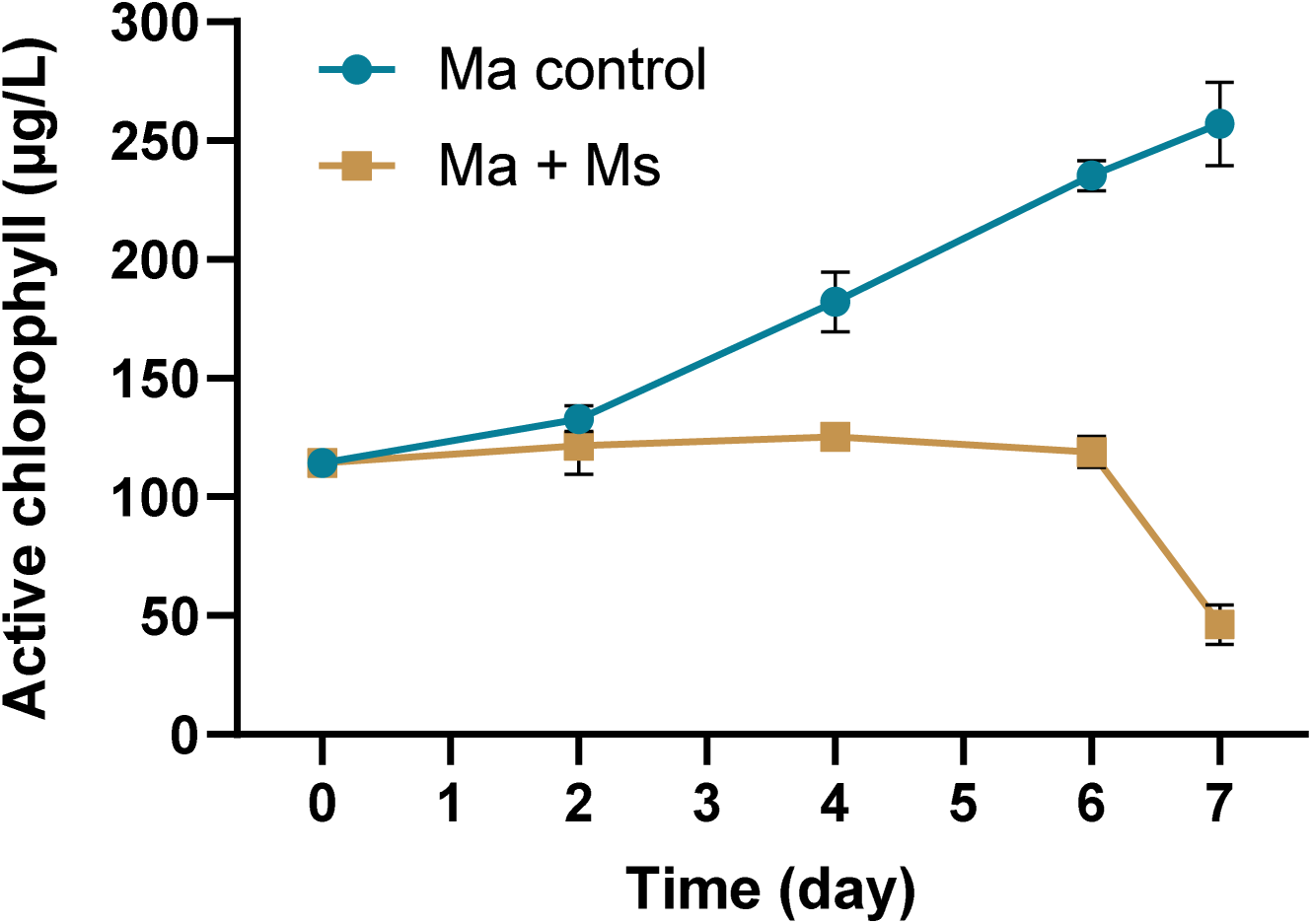
Growth comparison of Ma between monoculture and co-culture with Ms. Bars represent mean ± SD.

**Figure 4.**
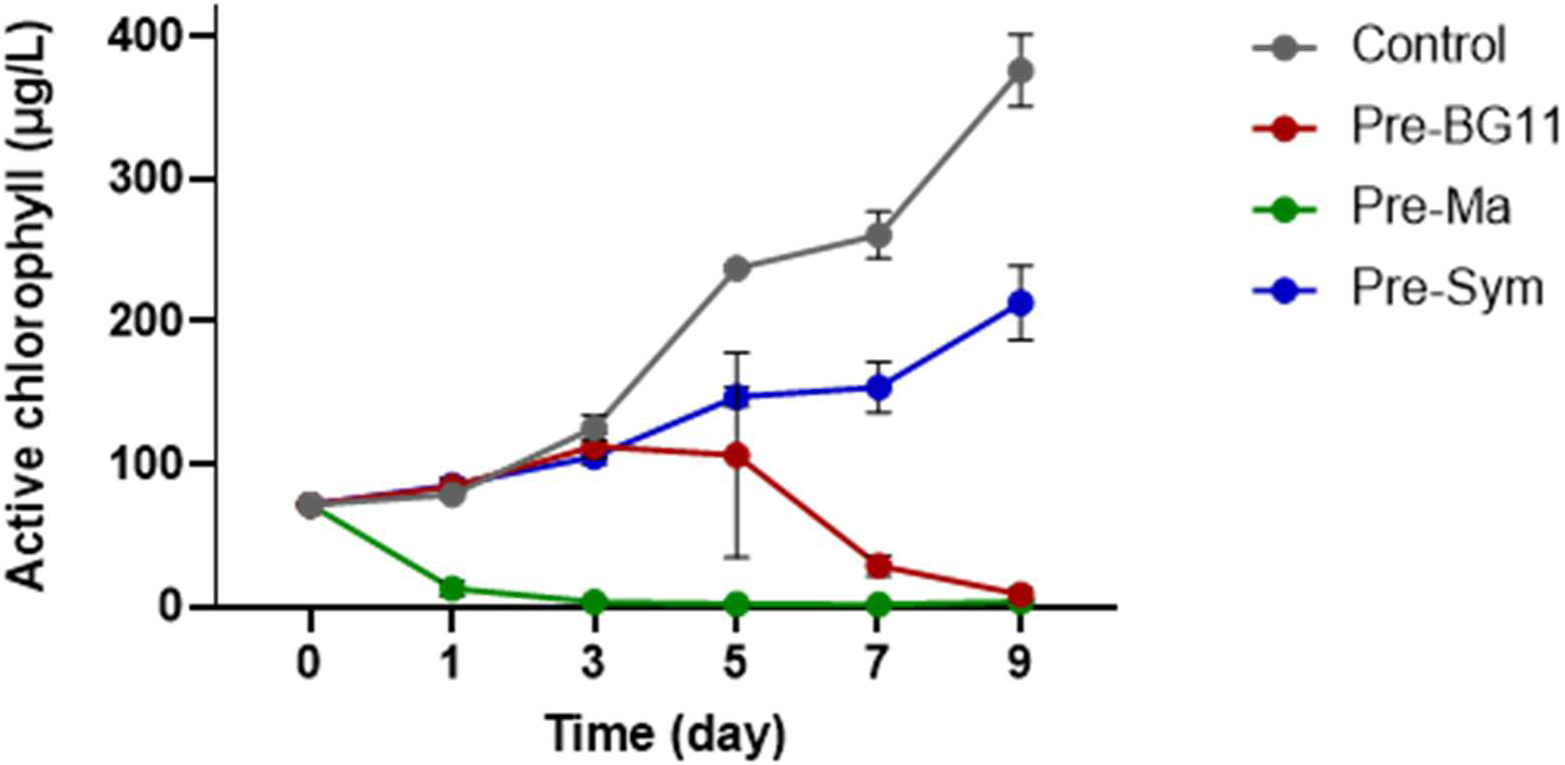
Growth of Ma in the co-culture with Ms pre-cultured under three distinct conditions (mean ± SD). Pre-BG11 (Ms pre-cultured in BG11 medium), Pre-Ma (Ms pre-cultured in Ma culture), and Pre-Sym (Ms pre-cultured in symbiotic microbiome of Ma).

### 3.2. Comparison of inhibitory effects of plant-derived chemicals and microbiome of Myriophyllum spicatum

Significant differences in inhibition were observed among the three groups on both experimental days (one-way ANOVA: day 1, F = 236.2, *p* = 1.97 × 10^-6^; day 2, F = 311.4, *p* = 8.69 × 10^-7^) (Figs. 2B and S2). On day 1, Group S exhibited a mean inhibition of 80%, which was significantly higher than that of both Group F (33%) (*p* = 2.7 × 10^-6^) and M (37%) (*p* = 4.2 × 10^-6^), while Group F and M did not differ significantly from each other (*p* = 0.377, Tukey’s HSD). By day 2, inhibition in Group F declined markedly to 3%, whereas Group M increased to 79% (*p* = 2.4 × 10^-6^). Group S showed the highest inhibition (91%), significantly exceeding Group M (*p* = 4.4 × 10^-2^) and F (*p* = 1.2 × 10^-6^).

### 3.3. Microbial community shifts linked to *Microcystis* inhibition

The ten most dominant bacterial genera in the co-culture microbial communities were *Microcystis*, *Pseudorhodobacter*, *Rhodobacter*, *Sphingopyxis*, *Pseudoxanthobacter*, *Devosia*, *Mesorthizobium*, *Roseococcus*, *Ferrovibrio*, and *Sphingorhabdus* (Fig. 5A). Among these, *Microcystis* was the most abundant genus, maintaining a substantial presence from inoculation until the onset of inhibition in each sample. While *Microcystis* remained abundant in the control, its relative abundance in Group Pre-Ma declined sharply throughout the experimental period. In Group Pre-BG11, a pronounced decrease was observed on day 7. Interestingly, *Microcystis* exhibited an increase in relative abundance in Group Pre-Sym. Several key genera demonstrated dynamic shifts in composition, particularly during *Microcystis* inhibition: *Pseudorhodobacter*, *Rhodobacter*, and *Sphingopyxis* declined with the reduction of *Microcystis*, but *Pseudoxanthobacter*, *Devosia*, *Mesorhizobium*, *Ferrovibrio*, *Pirellula*, and *Gemmatimonas* increased in relative abundance.

**Figure 5.**
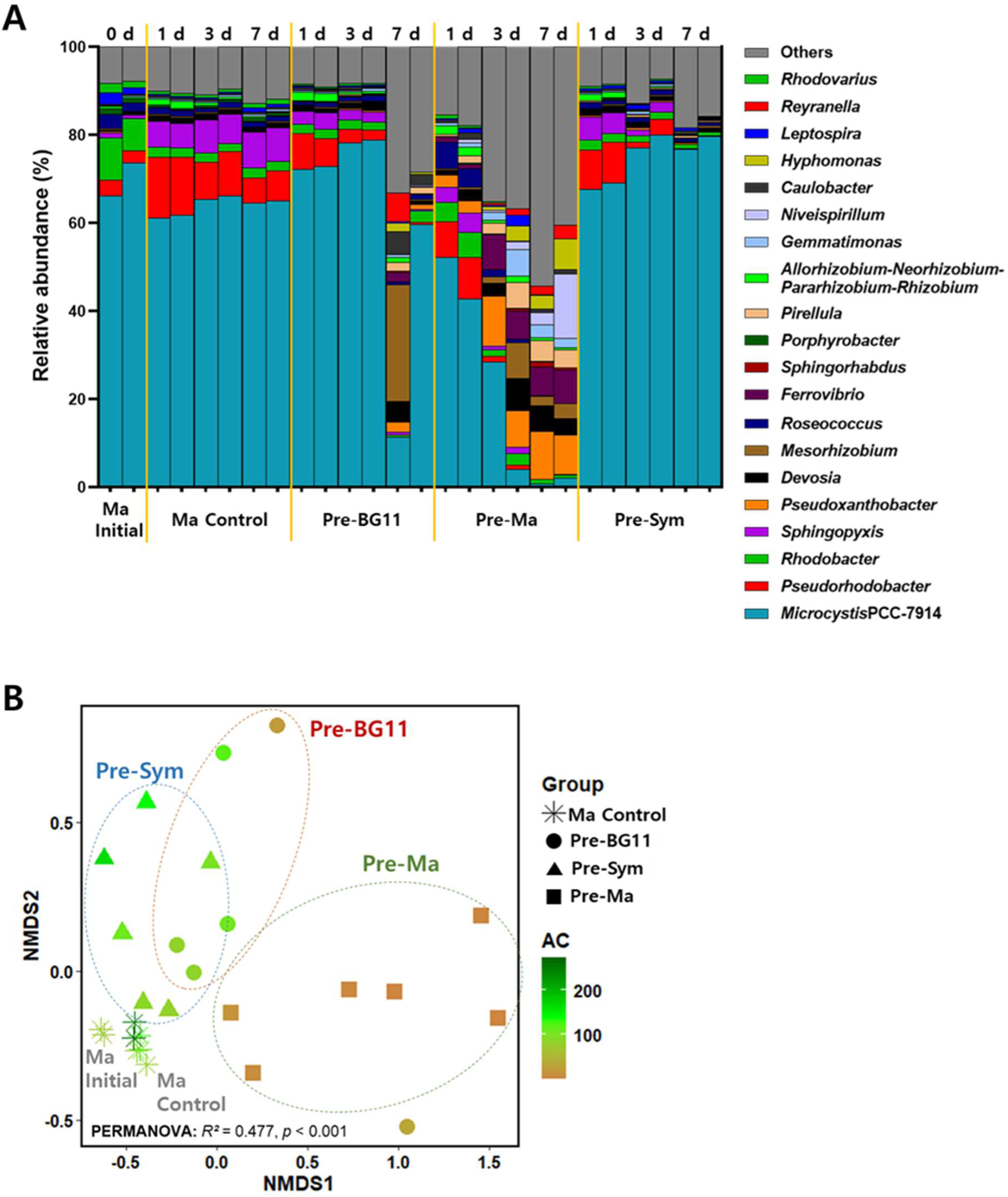
Shifts in microbial community composition in the Ma-Ms co-culture experiment across experimental conditions (Ma Initial, Ma Control, Pre-BG11, Pre-Ma, and Pre-Sym). (**A**) Genus-level community composition. (**B**) Non-metric multidimensional scaling (NMDS) ordination of microbial communities at the ASV level, based on Bray-Curtis dissimilarity (stress = 0.110). Each point represents an individual sample, with different shapes corresponding to experimental groups. The colour gradient represents active chlorophyll (AC) levels, ranging from tan (low) to dark green (high). Dotted ellipses indicate distinct clustering patterns among groups. Microbial community composition was significantly influenced by Group, Time, and AC, collectively explaining 70.4% of the total variation (PERMANOVA; Group: *R*^2^ = 0.477, F = 9.666, *p* < 0.001; Time: *R*^2^ = 0.151, F = 3.070, *p* = 0.032; AC: *R*^2^ = 0.076, F = 4.603, *p* = 0.023; 999 permutations).

Non-metric multidimensional scaling (NMDS) ordination of microbial communities at the ASV level revealed distinct clustering patterns among groups (Fig. 5B), indicating substantial shifts in community composition associated with *Microcystis* inhibition. Groups Pre-BG11 and Pre-Ma exhibited microbial assemblages that diverged from the control, with Group Pre-Ma showing the most pronounced shift, corresponding to the strongest inhibitory effect on *Microcystis*.

### 3.4. Key bacterial taxa related to *Microcystis* inhibition

Random forest analysis achieved a predictive accuracy over 97%, with the highest accuracy of 98% obtained using 25 features (Fig. S3). The area under the curve (AUC) reached 1.0 in the models using more than 10 variables (Fig. 6A), and the classes I and N were separated clearly (Fig. 6B). The average importance of the top-ranked features, which included bacterial taxa associated with *Microcystis* inhibition (red-colored in class I), is presented in Fig. 6C. A total of 28 bacterial taxa closely linked to the inhibitory response were identified (Fig. 6D), including *Bosea massiliensis*, *Pseudoxanthobacter* sp., *Pirellula* sp., *Ferrovibrio* sp., *Mesorhizobium* sp., *Reyranella aquatilis*, *Sphingomonas* sp., *Devosia insulae*, *Candidatus* Megaira sp., and *Bdellovibrio bacteriovorus*. The relative abundance of these bacteria was generally higher in *Microcystis*-inhibited group (I) compared to non-inhibited group (N) (Fig. S4). Several unclassified ASVs (e.g., ASV002, ASV005 and ASV019) were also detected (Table S1).

**Figure 6.**
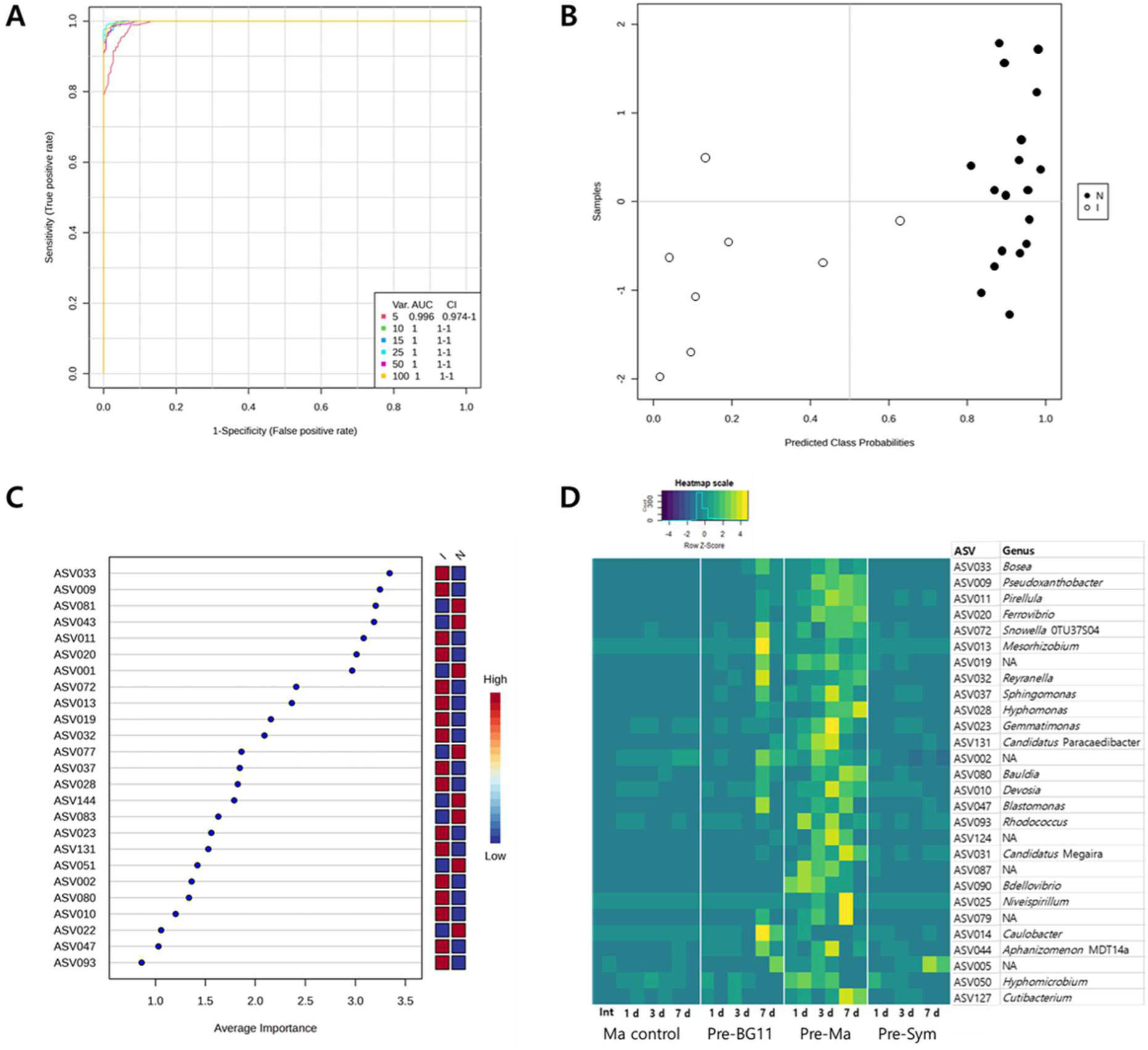
Random forest analysis of bacterial taxa associated with Ma inhibition. (**A**) Receiver operating characteristic (ROC) curves evaluating model performance in predicting inhibition levels. (**B**) Predicted class probabilities for samples categorized based on Ma inhibition (> 50%, I; < 50%, N). (**C**) Feature importance ranking of key bacterial taxa (ASVs) contributing to Ma inhibition, based on random forest regression. (**D**) Heatmap representing the Z-score transformed relative abundances of key bacterial taxa across experimental conditions over time.

### 3.5. Meta-transcriptomic analysis of microbial functions associated with *Microcystis* inhibition

Functional analysis at subsystem level 3 revealed distinct differences between the control (Ma monoculture) and the treatment (co-culture of Ma and Ms after > 50% inhibition of Ma). In the control group, Photosystem I, Photosystem II, and Phycobilisome, were predominant (Fig. 7A). In contrast, the treatment group exhibited significantly higher activities of β-Fimbriae, σ-Fimbriae, (GlcNAc)_2_ Catabolic Operon, and 5-FCL-like Protein (Fig. 7B). Comparison of functional activity levels further highlighted significant differences between the control and treatment groups. Photosynthesis-related functions (Photosystem I, Photosystem II, Phycobilisome, and Chlorophyll Biosynthesis), along with the Cyanobacterial Circadian Clock and Polyunsaturated Fatty Acids Synthesis, were significantly enriched in the control group (Fig. 7C). Conversely, bacterial adhesion-related functions (β-Fimbriae and σ-Fimbriae), carbohydrate metabolism-associated pathways ((GlcNAc)_2_ Catabolic Operon and 5-FCL-like Protein), ABC Transporter Branched-chain Amino Acid, and Bacterial Chemotaxis were notably elevated in the treatment group, where *Microcystis* growth was largely inhibited (Fig. 7D).

**Figure 7.**
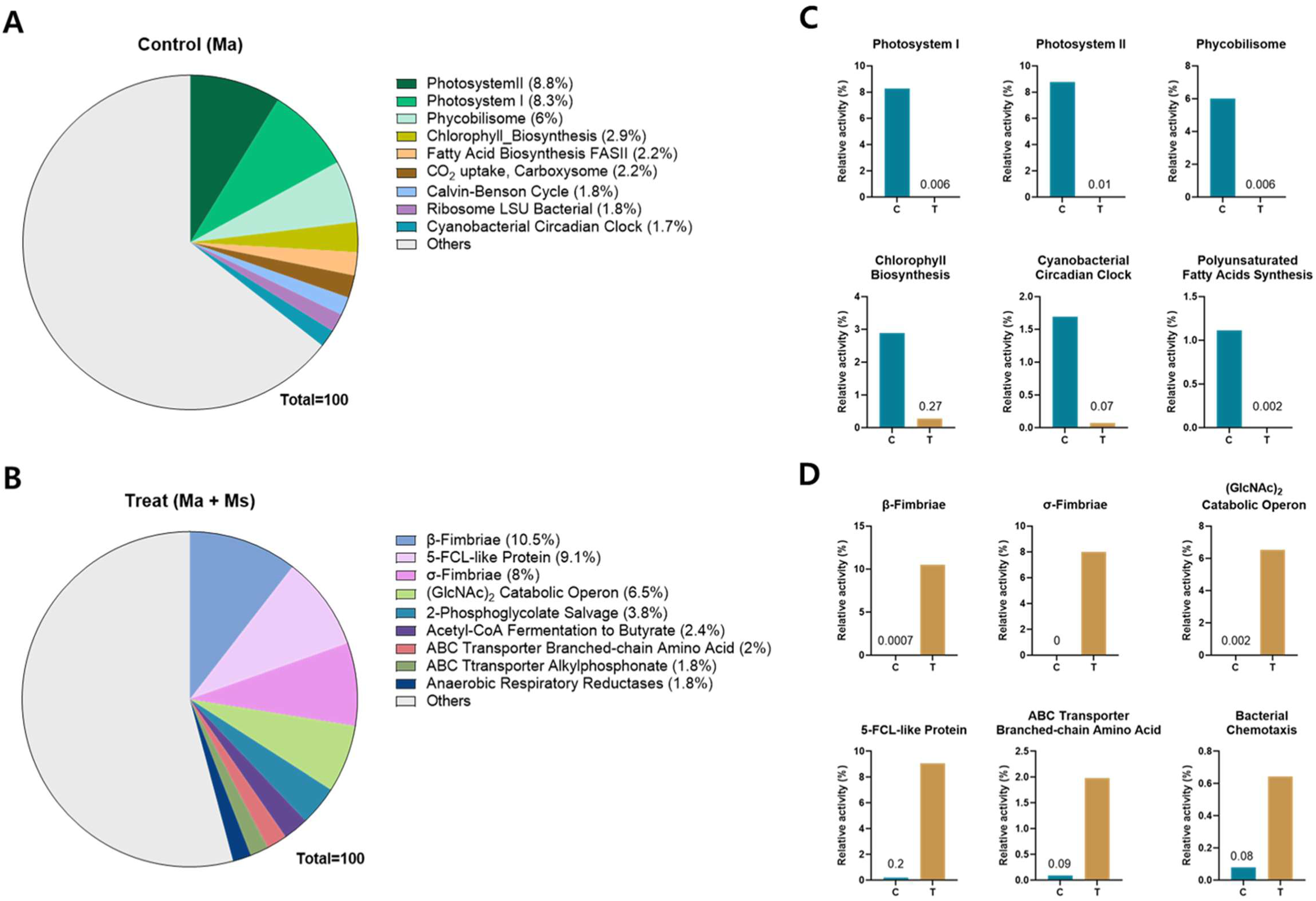
Meta-transcriptomic subsystem category distribution at hierarchy level 3 for control (**A**, Ma monoculture) and treatment (**B**, co-culture of Ma and Ms after > 50% inhibition of Ma). Comparison of relative activity levels of selected key functions between the control (**C**) and treatment (**D**).

## 4. Discussion

### 4.1. *Microcystis*-triggered inhibitory response

To determine whether Ma inhibition by Ms is constitutive or inducible, a pre-culture adaptation stage was implemented under three conditions: Pre-BG11, Pre-Ma, and Pre-Sym. When Ms pre-exposed to Ma (Pre-Ma) was subsequently re-exposed to Ma, it exhibited a rapid inhibitory response within the first day, suggesting that such prompt inhibitory effect is triggered by prior exposure to Ma (Figs. 1 and 4). In contrast, Group Pre-BG11 displayed a delayed onset of inhibition, with significant suppression observed on day 7, implying a potential lag phase in the algicidal response. This pattern closely resembles the onset of inhibition observed during the pre-culture phase of Pre-Ma (Fig. 3), indicating that the inhibitory response is not constitutive but rather triggered by exposure to Ma. Once established, the inhibitory response remained highly effective, as repeated inoculation of fresh Ma culture into the system resulted in rapid inhibition within 1 day. These findings are consistent with previous studies (Jeong et al., 2021; 2024), which demonstrated that daily re-inoculation of fresh Ma at the same concentration led to > 90% inhibition following initial exposure. Together, these results underscore the robust nature of the inhibitory mechanism, which, once established, maintains its effectiveness against recurring Ma exposure.

### 4.2. Comparison of inhibitory effects of plant-derived chemicals and microbiome of Myriophyllum spicatum

This study evaluated the inhibitory effects of three treatment groups: F (bacteria-free filtrate of the co-culture, representing plant-derived chemical effects only), S (culture solution containing both plant-derived chemicals and Ms*-*symbiotic microbiome), and M (Ms*-*symbiotic microbiome only). Among these, Group S showed the most rapid and potent inhibition, with significantly higher suppression of Ma on day 1 compared to Group F and M (Fig. 2B). By day 2, the inhibition in Group F had declined to negligible levels, suggesting that plant-derived chemical effects alone played a minimal role. In contrast, Group M exhibited inhibition exceeding 70% by day 2, while Group S showed an even greater inhibition than on day 1, further emphasizing the significant contribution of microbiome. Furthermore, repeated cycles of Ma-triggered inhibition promoted the maturation of biofilm on the inner surface of the experimental flask, and subsequent re-exposure of the matured biofilm to Ma resulted in rapid inhibition, further confirming the biofilm-mediated inhibitory effect (Fig. S5). These results suggest that microbiome-mediated interactions, rather than plant-derived chemical effects, are the primary drivers of Ma inhibition.

Until recently, the inhibitory effects of Ms on cyanobacterial growth have largely been attributed to its allelopathic activity (Jeong et al., 2021; Leu et al., 2002; Mohamed et al., 2017). However, the present study suggests that the plant-symbiotic microbiome plays a more dominant role in Ma inhibition, while the inhibitory effect of plant-derived allelochemicals appears to be minimal. This novel perspective aligns with recent advances in sequencing technology, which have provided deeper insights into the functional roles and ecological significance of plant-symbiotic microbiomes. Moreover, accumulating evidence on algicidal bacteria isolated from aquatic plants further supports these findings (Chen et al., 2023; Imai et al., 2021; Jiang et al., 2019; Miyashita et al., 2019). In addition, the inhibitory effect of Ms on Ma was consistently observed across different seasons (Fig. S6), and comparable effects were detected in several other freshwater submerged macrophytes (Fig. S7), suggesting that this phenomenon may extend beyond Ms. These results provide new insights into the potentially universal mechanisms underlying macrophyte-microbiome-mediated suppression of cyanobacterial growth.

### 4.3. Key bacterial taxa associated with *Microcystis* inhibition

Microbial community analysis identified several bacterial taxa significantly correlated with the inhibition of Ma. The dominant taxa included *Bosea massiliensis*, *Pseudoxanthobacter* sp., *Pirellula* sp., *Reyranella aquatilis*, *Sphingomonas* sp., *Hyphomonas* sp., *Gemmatimonas* sp., *Devosia insulae*, *Niveispirillum* sp., *Caulobacter daechungensis*, *Hyphomicrobium* sp., and *Cutibacterium* sp. (Fig. 6D and Table S1). Many of these taxa are recognized for their ability to degrade organic compounds, highlighting their potential contribution to nutrient cycling in the ecosystem (Huy et al., 2014; Liu et al., 2016; Pagnier et al., 2011; Yoon et al., 2014).

Notably, *Devosia* is well known for its exceptional capacity to degrade various toxins and its applications in bioremediation (Talwar et al., 2020), while *Rhodococcus* sp. can degrade organic pollutants, including petroleum, making it effective in bioremediation (Kim et al., 2018). Other taxa with adhesive properties, such as *Sphingomonas* sp. (Habimana et al., 2014; Bereschenko et al., 2010) and *Hyphomonas* sp. (Quintero and Weiner, 1995), are likely to contribute to biofilm formation, which may facilitate plant-microbiome interactions. Additionally, *Ferrovibrio* sp., involved in iron metabolism (Sorokina et al., 2012), and *Mesorhizobium* sp., known for nitrogen fixation and nutrient provision to plants (Verma et al., 2012; Yamaya-Ito et al., 2018), were also identified. Furthermore, some taxa, such as *Candidatus* Paracaedibacter, form symbiotic relationships with other microorganisms, indicating potential ecological interactions within the microbiome (Midha et al., 2021).

*Bdellovibrio bacteriovorus*, a predatory bacterium that invades and lyses Gram-negative bacteria, was identified as being associated with *Microcystis* inhibition (Fig. 6D and Table S1). It thrives in environments with high bacterial density, and exclusively preys on Gram-negative bacteria without exhibiting pathogenicity (Laloux, 2020; Lai et al., 2023). This taxon was detected on days 1 and 3 in Group Pre-Ma, with particularly high abundance on day 1 with 84% *Microcystis* inhibition, following the inoculation of fresh *Microcystis*. Similarly, *Candidatus* Megaira, an intracellular parasitic bacterium known to form symbiosis with freshwater algae (Davison et al., 2023; Pasqualetti et al., 2020), was also observed and is likely released during the lysis of Ma cells. Its z-score-transformed relative abundance peaked on day 7 in Group Pre-Ma (Fig. 6D), with Ma inhibition up to 99%. The normalized relative abundance of *Candidatus* Megaira increased from 0.11 ± 0.02% (day 1) to 3.39 ± 1.4% (day 7) in Group Pre-Ma, whereas *B. bacteriovorus* decreased from 0.33 ± 0.07% (day 1) to 0.05 ± 0.07% (day 7). Moreover, *B. bacteriovorus* was not detected in other groups. Such low abundance may explain why these bacteria have previously remained undetected or inconspicuous. Despite their low relative abundance, both bacteria warrant further investigation, since their presence appears closely linked to Ma cell lysis. Particularly, *Bdellovibrio* ranked as the 18th most abundant genus in the endophytic microbiota of Ms prior to co-cultivation (Figs. S8 and S9). This suggests that *Bdellovibrio* sp., originally residing as a plant symbiont, may have migrated out upon Ma inoculation in search of new prey. These findings highlight its potential cyanobactericidal role of predatory and symbiotic bacteria in Ma suppression and underscore the need for targeted approaches (e.g., specific primers) to enable their precise detection and quantification.

### 4.4. Putative inhibitory mechanisms based on meta-transcriptomic analysis

Functional analysis revealed distinct metabolic shifts in the microbial community following Ms-mediated inhibition of Ma. In the control group, photosynthesis-related functions, including Photosystem I, Photosystem II, Phycobilisome, and Chlorophyll Biosynthesis, were predominant, indicating active photosynthetic metabolism (Fig. 7A and C). Additionally, the enrichment of the Cyanobacterial Circadian Clock and Polyunsaturated Fatty Acid Synthesis suggests that Ma maintained stable physiological and regulatory processes under monoculture condition.

In contrast, the treatment group exhibited a significant reduction in photosynthesis-related functions, accompanied by a marked increase in bacterial adhesion-associated pathways (β-Fimbriae and σ-Fimbriae) and heterotrophic metabolic processes ((GlcNAc)_2_ Catabolic Operon and 5-FCL-like Protein). These changes indicate a functional shift in the microbial community following Ma inhibition (Fig. 7B and D).

Beta- and sigma-fimbriae usher proteins, key functional modules at hierarchy level 4 of β- and σ-Fimbriae, are essential for bacterial surface adhesion and biofilm formation, as they facilitate the assembly of fimbriae via the chaperone-usher (CU) pathway (Cheng et al., 2022; Pakharukova et al., 2022; Waksman and Hultgren, 2009). This CU system is responsible for building adhesive protein structures on the outer membrane of Gram-negative bacteria. Additionally, the σ-fimbriae tip adhesin FimH (Level 4), located at the end of type 1 fimbriae, represents a second major functional component of the σ-Fimbriae (Waksman and Hultgren, 2009; Sauer et al., 2016). The enrichment of fimbriae-related proteins likely reflects biofilm development on the plant surfaces and flask inner walls, which may facilitate the release of bacterial cells into the surrounding solution through biofilm dispersal (Greenwich et al., 2023; Sauer et al., 2022). These features suggest that biofilm formation may occur during interactions with Ma, and/or that predatory bacteria may utilize fimbriae to adhere to the cell surface of Ma. Fimbriae are well documented in *Bdellovibrio* and like organisms (BALOs) including *B. bacteriovorus* based on their predatory nature (Avidan et al., 2017). In addition, among the important taxa implicated in Ma inhibition (Fig. 6D and Table S1), several biofilm-forming bacteria, such as *Sphingomonas* (Min et al., 2009) and *Hyphomonas* (Quintero and Weiner, 1995; El Beaino et al., 2018), are also known to possess fimbriae.

The enrichment of (GlcNAc)_2_ Catabolic Operon in the treatment group suggests increased microbial activity related to *N*-acetylglucosamine (GlcNAc) metabolism. GlcNAc dimer is a key structural component of Gram-negative bacterial cell wall, forming part of the lipopolysaccharide that links lipid A and the core oligosaccharide (Lam et al., 2011). This operon enables bacteria to degrade (GlcNAc)_2_ into usable carbon and nitrogen sources, reflecting enhanced microbial utilization of organic matter released from lysed Ma cells. Additionally, elevated (GlcNAc)_2_ catabolism could be linked to increased bacterial turnover and the degradation of cyanobacterial biomass. The predatory bacterium *Bdellovibrio* is known to discriminate prey-associated lipopolysaccharide O-antigens, and to mediate recognition and adhesion to specific glycan epitopes, such as GlcNAc, thereby influencing predatory interactions (Caulton et al., 2024; Duncan et al., 2018). These observations raise the possibility that (GlcNAc)_2_ may be involved in mediating microbial interactions during Ma inhibition. Further research is required to determine whether (GlcNAc)_2_ functions as a chemoattractant for predatory bacteria that preferentially target Ma.

The upregulation of 5-FCL-like Protein, a key enzyme in folate metabolism, proposes increased bacterial involvement in one-carbon metabolism, a critical pathway for nucleotide biosynthesis and cellular growth. These metabolic shifts imply that microbial communities in the treatment group adapted to the altered nutrient environment by increasing their ability to degrade and assimilate organic compounds. The disintegration of Ma cells results in the release of organic matter into the environment, thereby promoting bacterial carbohydrate metabolism. The enrichment of 5-FCL-like Protein suggests an increased demand for folate-dependent reactions, potentially driven by elevated bacterial proliferation and metabolic activity (Li et al., 2021). These findings point to a transition toward enhanced microbial decomposition and resource utilization, supporting a shift toward heterotrophic metabolism within the microbial community.

The enrichment of ABC Transporter Branched-chain Amino Acid (BCAA) metabolism in the treatment group suggests increased microbial competition for amino acids (Gesbert et al., 2015), following Ma inhibition. BCAAs (valine, leucine, and isoleucine) are essential for bacterial growth and metabolism, and their active transport indicates a shift in nutrient acquisition dynamics. The suppression of Ma likely altered the availability of organic nitrogen, prompting heterotrophic bacteria to upregulate BCAA transport for survival and metabolic adaptation (Dutta et al., 2022; Kaiser and Heinrichs, 2018).

Additionally, the increased abundance of Bacterial Chemotaxis functions suggests that bacteria actively migrated toward newly available nutrient sources in the treatment groups (Colin et al., 2021). Chemotaxis allows bacteria to sense and respond to chemical gradients, facilitating colonization in new favorable niches and optimizing resource utilization. This response may also contribute to biofilm formation through crosstalk between chemotaxis and biofilm formation pathways (Huang et al., 2019; Oliveira et al., 2016; Wei and Ma, 2013). Considering that biofilm formation and fimbrial adhesion are typical characteristics of infectious or predatory bacteria, the observed features suggest that similar strategies may be involved in Ma inhibition, potentially by promoting microbial aggregation.

These findings of meta-transcriptome study indicate that Ma inhibition is primarily driven by the Ms-associated microbial community. This microbiome promotes both structural and functional reconfiguration of the system, shifting it from a cyanobacteria-dominated state to one characterized by enhanced heterotrophic activity and bacterial adhesion processes. More specifically, Ma inhibition appears to be mediated by coordinated microbiome-driven processes, including biofilm formation, organic matter recycling, and metabolic reprogramming, which together contribute to sustained suppression of cyanobacterial growth.

Furthermore, two bacterial strains were identified from the co-culture of Ms and Ma that are closely associated with Ma inhibition. These results underscore the intricate ecological interactions among macrophytes, their symbiotic microbiomes, and cyanobacteria, highlighting the potential involvement of microbiome-mediated cyanobactericidal effects. This study advances the understanding of macrophyte-cyanobacteria interactions and their broader ecological implications, while emphasizing the need for further investigation into the specific inhibitory mechanisms and metabolic pathways underlying these complex interspecies dynamics.

### 4.5. Potential for *Microcystis* bloom control as NbS

Recent studies have increasingly explored selective and eco-friendly NbS for controlling cyanoHABs without causing secondary ecological disturbances. Freshwater macrophytes offer a promising approach for improving water quality while minimizing ecosystem disruption. Various aquatic plants, such as *Trapa* (Miyashita et al., 2019; Akao et al., 2014), *Potamogeton* (Vanderstukken et al., 2011), *Vallisneria* (Jiang et al., 2019; Maredová et al., 2021), and *Myriophyllum* (Švanys et al., 2014; Gao et al., 2022), have been investigated for their potential to control Ma blooms. However, many of these studies rely on plant extracts, which often require high concentrations, posing challenges for practical field application. In contrast, our findings demonstrate that Ma suppression by Ms is primarily driven by its symbiotic microbiome rather than plant-derived allelopathic compounds. Moreover, the observed shifts in bacterial community composition following Ma inhibition suggest a potential microbial succession that may reinforce bloom suppression. Furthermore, Ms effectively suppressed a natural *Microcystis* bloom sample collected from Daecheong Reservoir, South Korea (Fig. S10), supporting the potential applicability of this approach under natural bloom conditions. This microbiome-driven process could provide a more sustainable and self-maintaining mechanism for bloom control compared to extract-based approaches.

The proposed strategy, which utilizes submerged plants in conjunction with their symbiotic microbiomes (biofilms), offers several advantages over conventional methods. This approach not only enhances carbon sequestration but also minimizes ecological disturbance, promotes biodiversity, and improves overall ecosystem health. In addition, its ability to remove excess nutrients such as phosphorus and nitrogen highlights its potential as a viable NbS model for sustainable water quality management.

Further research should focus on isolating and characterizing the functional roles of key bacterial taxa identified in this study, as well as evaluating their effectiveness under natural freshwater conditions. In particular, identifying functionally relevant microbial consortia and validating their performance at the ecosystem scale will be essential for translating these findings into practical bloom management strategies. These efforts will be critical for developing scalable, microbiome-based solutions for long-term cyanobacterial bloom control.

## 5. Conclusion

This study demonstrates that the microbiome associated with *Myriophyllum spicatum* plays a more significant role in inhibiting *Microcystis aeruginosa* compared to the plant-derived allelochemicals. The inhibitory response, triggered by exposure to *Microcystis aeruginosa*, was accompanied by shifts in the microbial community composition, leading to faster inhibition upon re-exposure. Key bacterial taxa involved in *Microcystis* inhibition showed functional traits related to adhesion, biofilm formation, and predation, highlighting diverse microbial characteristics observed during the *Microcystis* suppression. Many of these taxa are also known to possess the capability to degrade organic matter, suggesting a potential role in the utilization of organic material released during *Microcystis* decline. These findings underscore the critical role of aquatic plant-symbiotic microbiomes in cyanobacterial suppression. Leveraging such microbiome-plant interactions presents a promising and eco-friendly NbS for mitigating cyanoHABs, warranting further application-oriented and field-based studies.

## Acknowledgments

This research was supported by National Institute of Environmental Research (NIER) (NIER-2024-04-02-084), Korea Environmental Industry & Technology Institute (KEITI) through Aquatic Ecosystem Conservation Research Program (2022003050004), the National Research Council of Science & Technology (NST) grant by the Korea government (MSIT) (No. GTL25021-310), and Korea Research Institute of Bioscience and Biotechnology (KRIBB) Research Initiative Program (KGM1242612). We are grateful to senior and fellow researchers for their thoughtful interest and insightful questions that enriched this work.

## Author contributions

**Seonah Jeong:** Conceptualization, Writing – original draft, Writing – review & editing, Resources, Methodology, Investigation, Formal analysis, Data curation, Software, Visualization, Project administration. **Hayoung Lee:** Writing – original draft, Resources, Investigation, Formal analysis, Visualization. **So-Ra Ko:** Methodology, Resources, Formal analysis, Project administration. **Dong-Yun Choi:** Resources, Formal analysis, Project administration. **Won-Suk Choi:** Methodology, Resources, Formal analysis. **Yuna Shin:** Writing – review & editing, Project administration. **Kyunghyun Kim:** Writing – review & editing, Project administration. **Hee-Sik Kim:** Writing – review & editing, Funding acquisition. **Chi-Yong Ahn:** Conceptualization, Writing – review & editing, Supervision, Project administration, Funding acquisition.

## Data availability

The raw sequencing data reported in this paper have been deposited in the Genome Sequence Archive (GSA) in National Genomics Data Center, China National Center for Bioinformation / Beijing Institute of Genomics, Chinese Academy of Sciences (GSA: CRA027591), and are publicly accessible at https://ngdc.cncb.ac.cn/gsa. The remaining data are available at https://ngdc.cncb.ac.cn/omix/release/OMIX010600 (accession no. OMIX010600) and https://ngdc.cncb.ac.cn/bioproject (accession no. PRJCA041695).

## List of abbreviations

ASV: Amplicon Sequence Variant
CU: Chaperone-Usher
Ma: Microcystis aeruginosa
Ms: Myriophyllum spicatum
MCCV: Monte-Carlo Cross Validation
NbS: Nature-based Solutions
NMDS: Non-metric Multidimensional Scaling
ROC: Receiver Operating Characteristic

