## supplementary for "*Microcystis*-triggered shifts in the symbiotic microbiome of *Myriophyllum spicatum* rapidly suppress *Microcystis aeruginosa*"

**Chi-Yong Ahn**

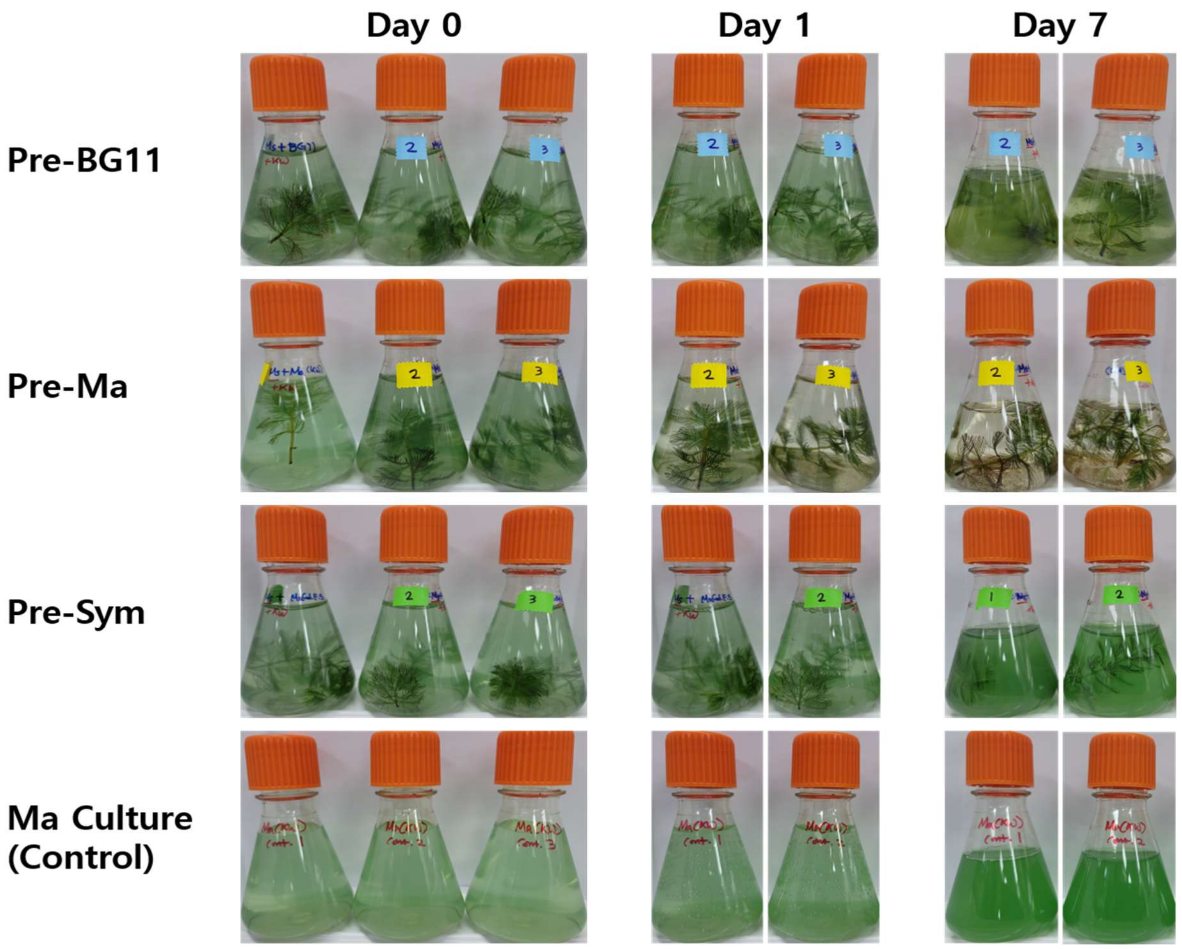


**Fig. S1.** Representative photographs of the experiments shown in Figure 1, the subsequent co-culture experiments following the pre-culture treatments and *Microcystis aeruginosa* (Ma) control group.

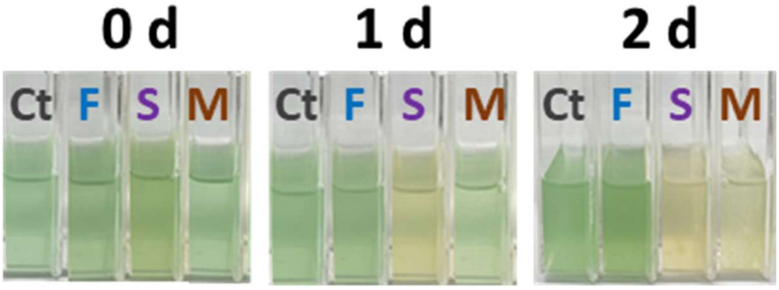


**Fig. S2.** Comparison of inhibitory effects of chemicals and microbiome of *Myriophyllum spicatum*: Ct, the control group with *Microcystis aeruginosa* only; F, bacteria-free filtrate of the co-culture, representing plant-derived chemical effects only; S, culture solution containing both plant-derived chemical substances and plant-symbiotic microbiomes; M, *Myriophyllum*-symbiotic microbiome effects only.

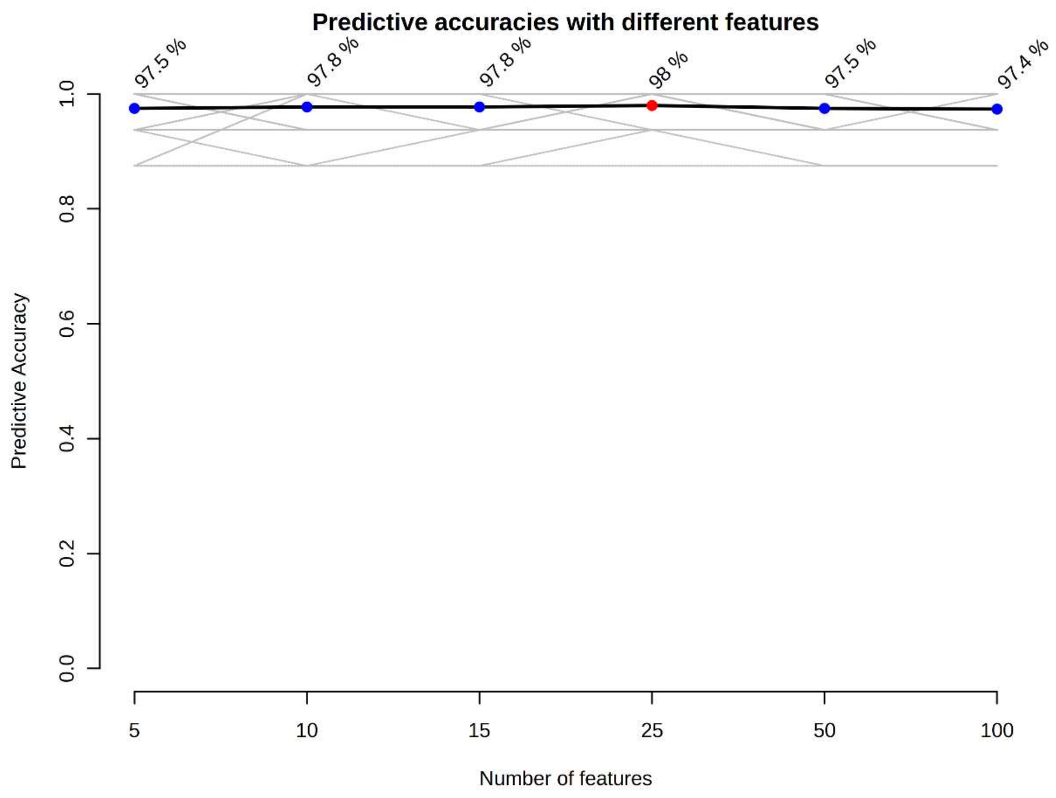


**Fig. S3.** The predicted accuracy of random forest model based on the number of features.

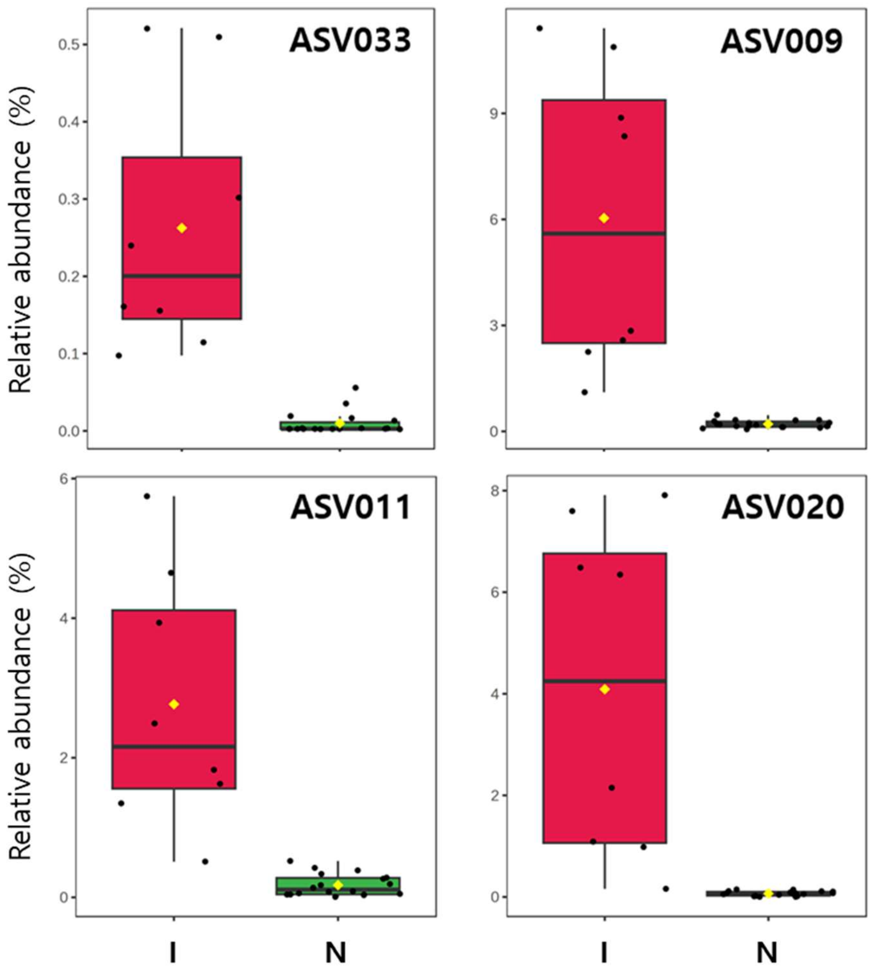


**Fig. S4.** Relative abundance of the top 4 key bacterial taxa based on inhibition level (I, inhibition > 50% on *Microcystis*; N, inhibition < 50%).

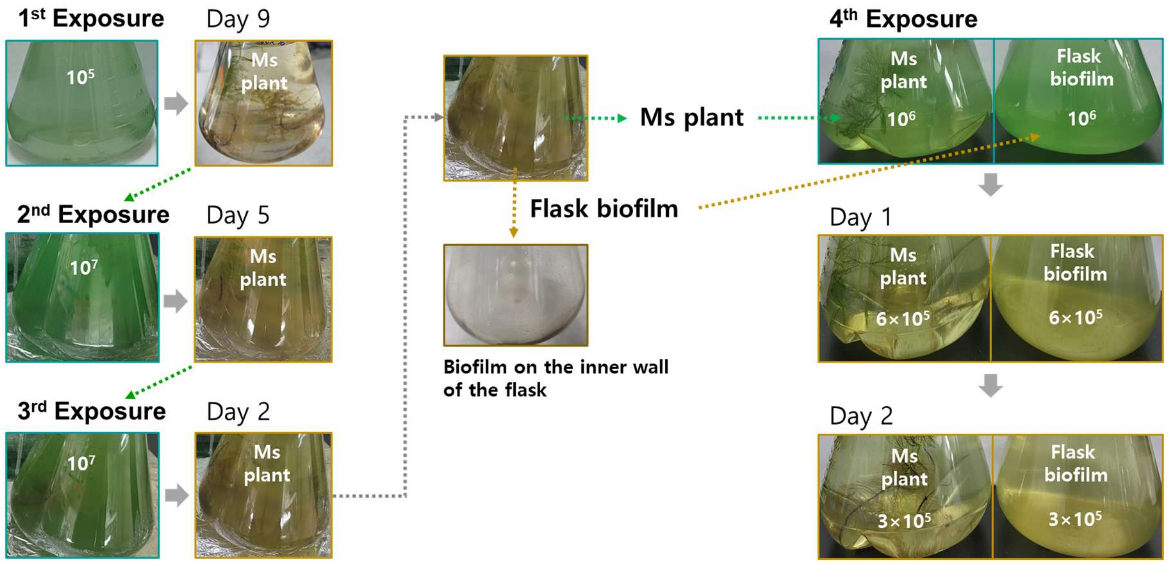


**Fig. S5.** Rapid inhibition of *Microcystis aeruginosa* (Ma) by flask-inner-surface biofilm formed during co-culture with *Myriophyllum spicatum* (Ms). Ma cultures (1 L) with initial cell density of $10^5 – 10^7$ cells/mL were co-cultivated with a 20-cm shoot of Ms. Following three consecutive cycles of Ma replenishment and subsequent growth inhibition, the culture solution was removed, leaving only the visible biofilm attached to the inner wall of the culture flask. A fresh Ma culture was then introduced into the biofilm-only flask and compared with a control flask containing Ms. Representative photographs show that both the Ms-containing flask and the biofilm-only flask reduced Ma cell density from approximately $10^6$ to $3×10^5$ cells/mL by day 2. The comparable inhibitory effect observed in the absence of the plant suggests that biofilm-associated microbial communities contribute substantially to the inhibition of Ma growth.

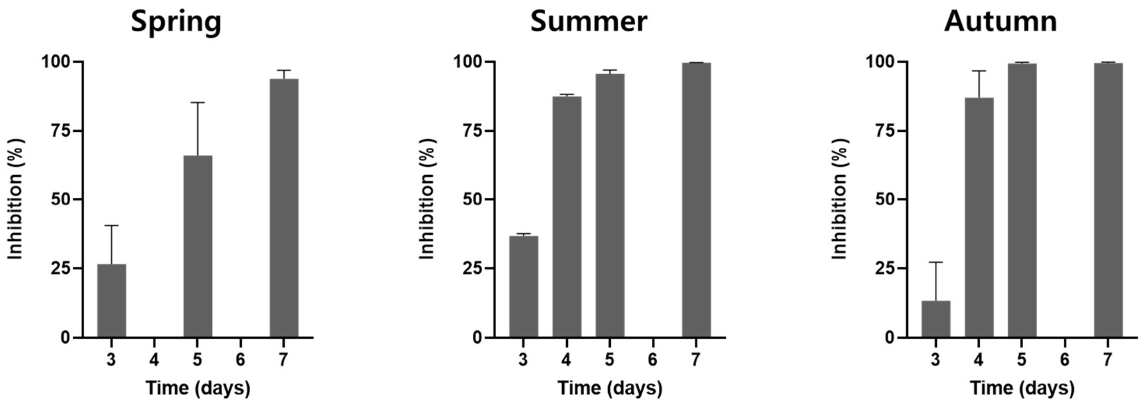


**Fig. S6.** Seasonal variation in the inhibitory effects of Ms on Ma. Ms plants were collected in May (Spring), July (Summer), and October (Autumn) 2024 from urban streams in Daejeon, South Korea. Apical shoots (5 cm) were co-cultured with Ma ($9.5 \times 10^5$ cells/mL) in BG11 medium. Inhibition was determined from active chlorophyll measurements and expressed relative to the control. Values represent mean ± SD ($n = 2$). Inhibition exceeded 50% on day 5 for spring-collected plants, and on day 4 for summer- and autumn-collected plants.

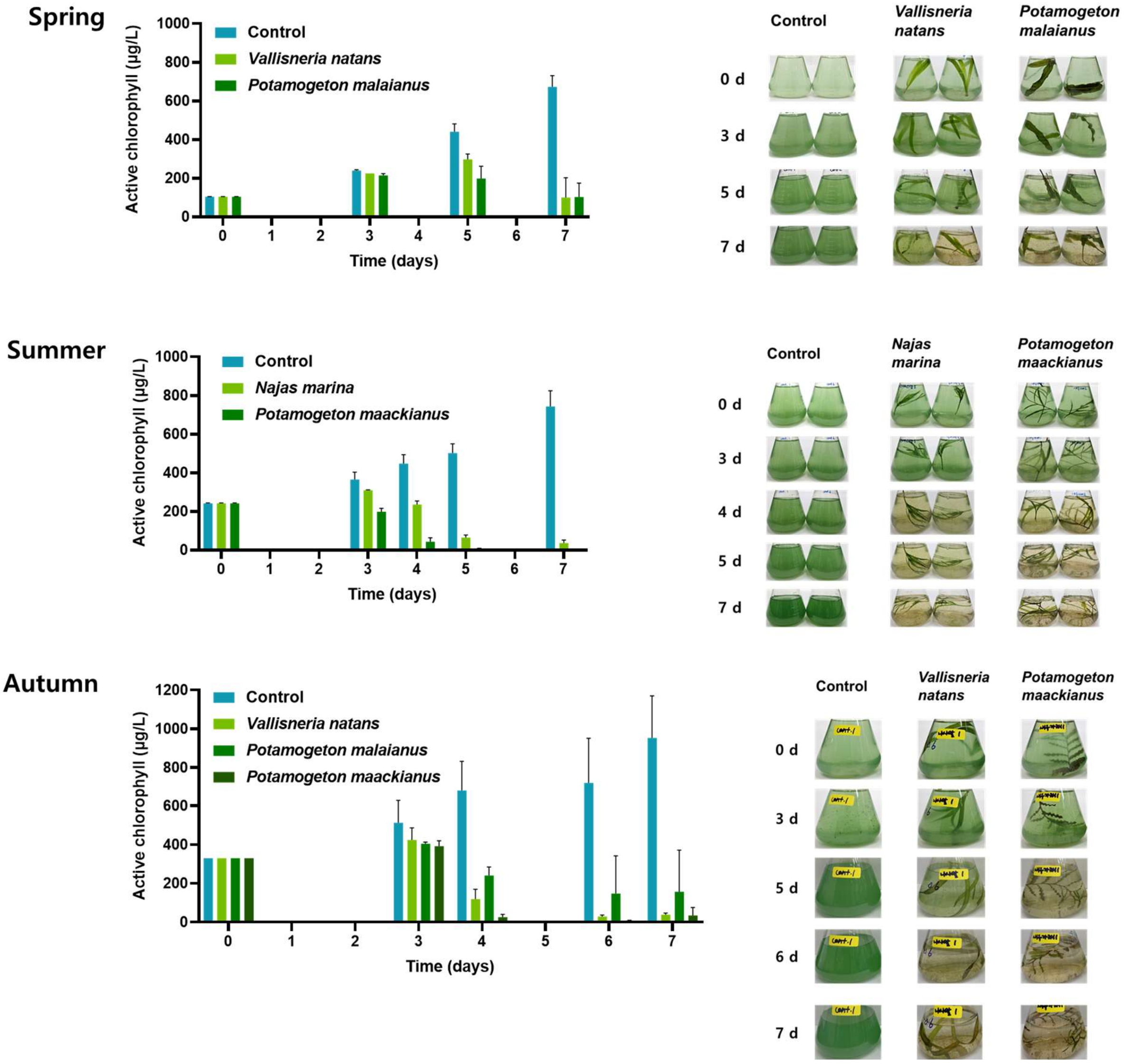


**Fig. S7.** Seasonal variation in the inhibitory effects of submerged freshwater macrophytes on the growth of Ma. Three submerged macrophyte species (*Vallisneria natans*, *Potamogeton malaianus*, and *Potamogeton maackianus*) were collected during Spring (May), Summer (July), and Autumn (October) in 2024, and co-cultured with Ma ($9.5 \times 10^5$ cells/mL) in BG11 medium. Changes in active chlorophyll level were monitored over the experimental period, and representative photographs were taken simultaneously to document visual changes in culture flasks. Data are presented as mean ± SD from two biological replicates.

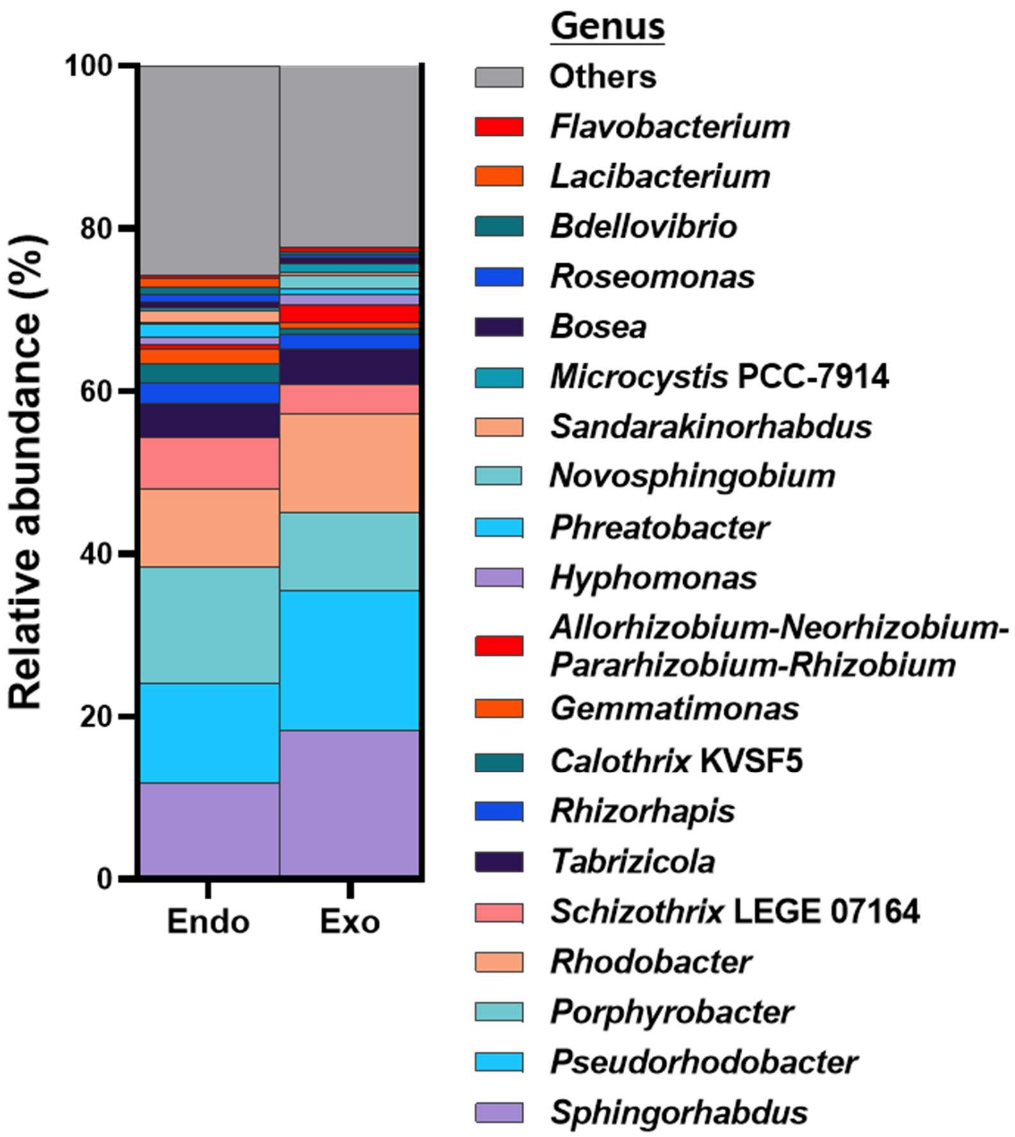


**Fig. S8.** Genus-level bacterial community composition of the endophytic (Endo) and exophytic (Exo) microbiomes of *Myriophyllum spicatum* collected from the Yudeung Stream, South Korea.

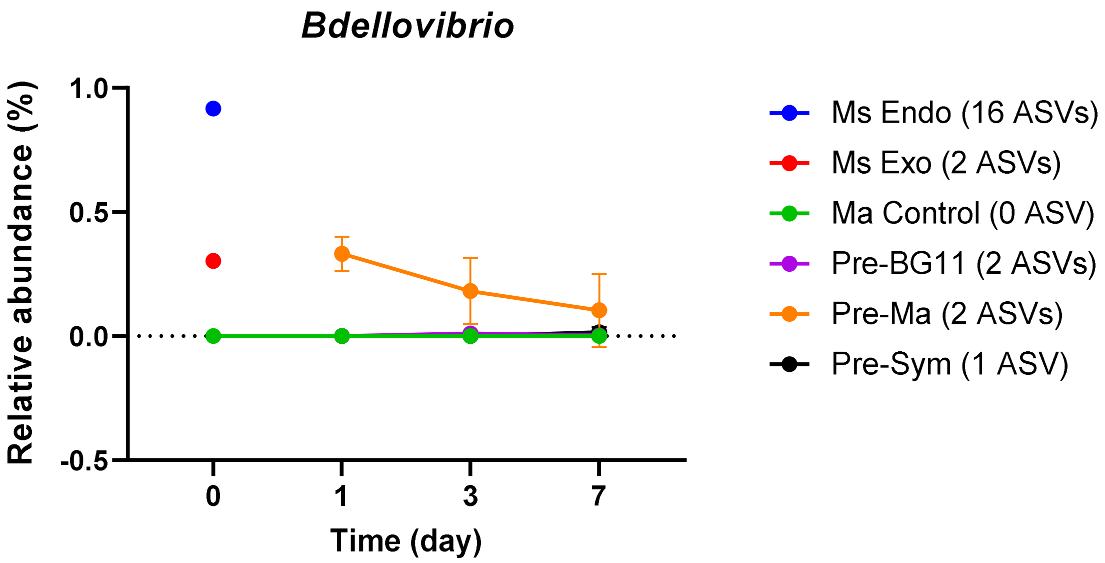


**Fig. S9.** Sum of relative abundance of total 20 *Bdellovibrio* sp. strains (ASVs).

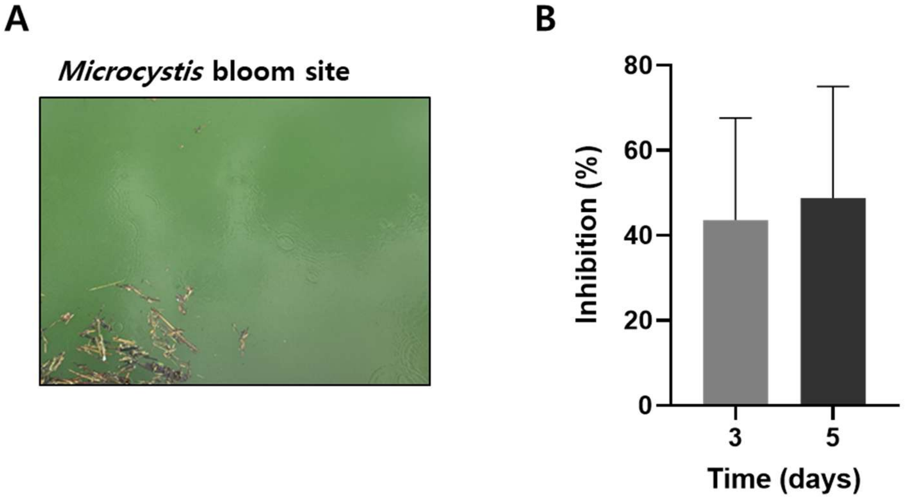


**Fig. S10.** Inhibitory effect of the Ms-associated microbiome on natural *Microcystis* bloom sample. Natural *Microcystis* bloom sample (2 L) was collected in August 2024 from Daecheong Reservoir, South Korea (A). The initial abundance of *Microcystis* spp. in the bloom sample was 2.6 × $10^4$ cells/mL. Two shoots of *M. spicatum* (each 20 cm in length) were added to the 20-L *Microcystis* bloom sample. The bar graph presents inhibition levels at day 3 and 5 (B). Data are presented as mean ± SD from two biological replicates.

**Table S1.** Key bacterial ASVs associated with *Microcystis aeruginosa* inhibition (corresponding to Fig. 7D), including their taxonomic classification from phylum to species.

| ASV | Frequency | Importance | Phylum | Class | Order | Family | Genus | Species |
|---|---|---|---|---|---|---|---|---|
| ASV033 | 0.88 | 3.34 | Proteobacteria | *Alphaproteobacteria* | *Rhizobiales* | *Beijerinckiaceae* | *Bosea* | *massiliensis* |
| ASV009 | 0.90 | 3.25 | Proteobacteria | *Alphaproteobacteria* | *Rhizobiales* | *Xanthobacteraceae* | *Pseudoxanthobacter* | NA |
| ASV011 | 0.80 | 3.08 | Planctomycetota | *Planctomycetes* | *Pirellulales* | *Pirellulaceae* | *Pirellula* | NA |
| ASV020 | 0.80 | 3.01 | Proteobacteria | *Alphaproteobacteria* | *Ferrovibrionales* | *Ferrovibrionaceae* | *Ferrovibrio* | NA |
| ASV072 | 0.42 | 2.41 | Cyanobacteria | *Cyanobacteriia* | *Cyanobacteriales* | *Microcystaceae* | *Snowella* 0TU37S04 | NA |
| ASV013 | 0.40 | 2.36 | Proteobacteria | *Alphaproteobacteria* | *Rhizobiales* | *Rhizobiaceae* | *Mesorhizobium* | NA |
| ASV019 | 0.28 | 2.16 | Proteobacteria | *Alphaproteobacteria* | SAR11 clade | Clade III | NA | NA |
| ASV032 | 0.40 | 2.09 | Proteobacteria | *Alphaproteobacteria* | *Reyranellales* | *Reyranellaceae* | *Reyranella* | *aquatilis* |
| ASV037 | 0.20 | 1.84 | Proteobacteria | *Alphaproteobacteria* | *Sphingomonadales* | *Sphingomonadaceae* | *Sphingomonas* | NA |
| ASV028 | 0.24 | 1.82 | Proteobacteria | *Alphaproteobacteria* | *Caulobacterales* | *Hyphomonadaceae* | *Hyphomonas* | NA |
| ASV023 | 0.18 | 1.56 | Gemmatimonadota | *Gemmatimonadetes* | *Gemmatimonadales* | *Gemmatimonadaceae* | *Gemmatimonas* | NA |
| ASV131 | 0.20 | 1.53 | Proteobacteria | *Alphaproteobacteria* | *Paracaedibacterales* | *Paracaedibacteraceae* | *Candidatus* Paracaedibacter | NA |
| ASV002 | 0.18 | 1.36 | Proteobacteria | *Alphaproteobacteria* | NA | NA | NA | NA |
| ASV080 | 0.08 | 1.34 | Proteobacteria | *Alphaproteobacteria* | *Rhizobiales* | *Rhizobiales Incertae Sedis* | *Bauldia* | NA |
| ASV010 | 0.10 | 1.20 | Proteobacteria | *Alphaproteobacteria* | *Rhizobiales* | *Devosiaceae* | *Devosia* | *insulae* |
| ASV047 | 0.10 | 1.03 | Proteobacteria | *Alphaproteobacteria* | *Sphingomonadales* | *Sphingomonadaceae* | *Blastomonas* | NA |
| ASV093 | 0.04 | 0.86 | Actinobacteriota | *Actinobacteria* | *Corynebacteriales* | *Nocardiaceae* | *Rhodococcus* | NA |
| ASV124 | 0.04 | 0.82 | Actinobacteriota | *Actinobacteria* | *Micrococcales* | *Microbacteriaceae* | NA | NA |
| ASV031 | 0.02 | 0.77 | Proteobacteria | *Alphaproteobacteria* | *Rickettsiales* | *Rickettsiaceae* | *Candidatus* Megaira | NA |
| ASV087 | 0.02 | 0.72 | Desulfobacterota | *Desulfuromonadia* | *Bradymonadales* | NA | NA | NA |
| ASV090 | 0.02 | 0.70 | Bdellovibrionota | *Bdellovibrionia* | *Bdellovibrionales* | *Bdellovibrionaceae* | *Bdellovibrio* | *bacteriovorus* |
| ASV025 | 0.04 | 0.66 | Proteobacteria | *Alphaproteobacteria* | *Azospirillales* | *Azospirillaceae* | *Niveispirillum* | NA |
| ASV079 | 0.02 | 0.58 | Proteobacteria | *Alphaproteobacteria* | *Sphingomonadales* | *Sphingomonadaceae* | NA | NA |
| ASV014 | 0.06 | 0.56 | Proteobacteria | *Alphaproteobacteria* | *Caulobacterales* | *Caulobacteraceae* | *Caulobacter* | *daechungensis* |
| ASV044 | 0.04 | 0.52 | Cyanobacteria | *Cyanobacteriia* | *Cyanobacteriales* | *Nostocaceae* | *Aphanizomenon* MDT14a | NA |
| ASV005 | 0.02 | 0.32 | Proteobacteria | *Alphaproteobacteria* | *Azospirillales* | *Azospirillaceae* | NA | NA |
| ASV050 | 0.02 | 0.15 | Proteobacteria | *Alphaproteobacteria* | *Rhizobiales* | *Hyphomicrobiaceae* | *Hyphomicrobium* | NA |
| ASV127 | 0.02 | 0.06 | Actinobacteriota | *Actinobacteria* | *Propionibacteriales* | *Propionibacteriaceae* | *Cutibacterium* | NA |
